# The Quartet Data Portal: integration of community-wide resources for multiomics quality control

**DOI:** 10.1101/2022.09.26.507202

**Authors:** Jingcheng Yang, Yaqing Liu, Jun Shang, Qiaochu Chen, Qingwang Chen, Luyao Ren, Naixin Zhang, Ying Yu, Zhihui Li, Yueqiang Song, Shengpeng Yang, Andreas Scherer, Weida Tong, Huixiao Hong, Leming Shi, Wenming Xiao, Yuanting Zheng

**Author notes:** These authors contributed equally: Jingcheng Yang, Yaqing Liu, Jun Shang. Corresponding author’s.

## Abstract

The implementation of quality control for multiomic data requires the widespread use of well-characterized reference materials, reference datasets, and related resources. The Quartet Data Portal was built to facilitate community access to such rich resources established in the Quartet Project. A convenient platform is provided for users to request the DNA, RNA, protein, and metabolite reference materials, as well as multi-level datasets generated across omics, platforms, labs, protocols, and batches. Interactive visualization tools are offered to assist users to gain a quick understanding of the reference datasets. Crucially, the Quartet Data Portal continuously collects, evaluates, and integrates the community-generated data of the distributed Quartet multiomic reference materials. In addition, the portal provides analysis pipelines to assess the quality of user-submitted multiomic data. Furthermore, the reference datasets, performance metrics, and analysis pipelines will be improved through periodic review and integration of multiomic data submitted by the community. Effective integration of the evolving technologies via active interactions with the community will help ensure the reliability of multiomics-based biological discoveries. The Quartet Data Portal is accessible at https://chinese-quartet.org.

**Graphical Abstract:** 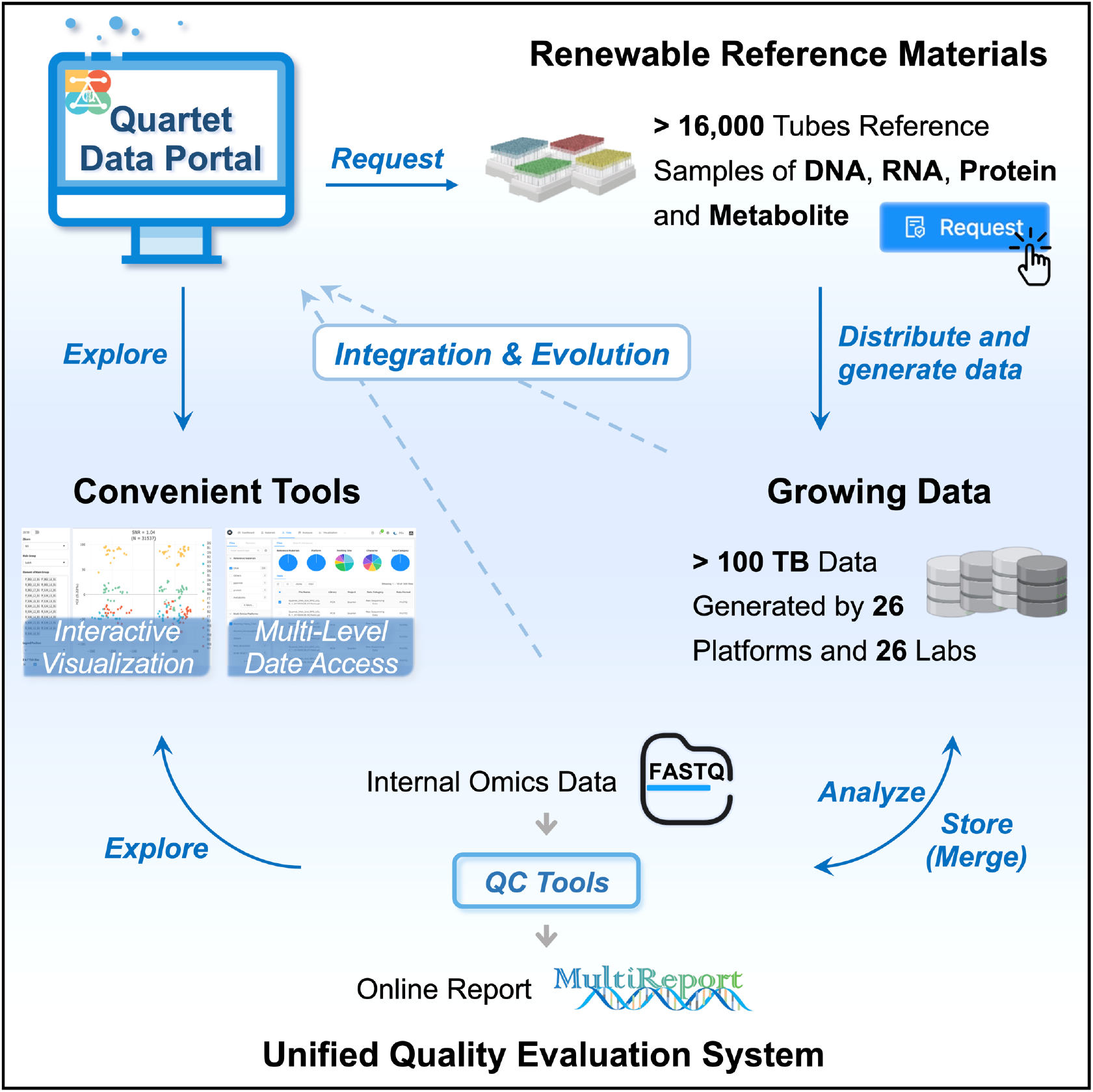

---

Well-characterized reference materials are powerful resources to promote the reliability of multiomic profiling, yet efforts to improve their usability by providing a one-stop solution are still in its infancy^1–3^. To date, no data portal has fully met the needs of users for integrated resources involving the entire chain from sample acquisition to data quality assessment. The general way to obtain the resources is limited and centralized, mainly through publications and official portals. The later often only provides links to various fragmented information, e.g., data, tools, and publications. Typical examples are the websites of MAQC/SEQC^4–7^, GIAB^8, 9^, HUPO-PSI^10^ and mQACC^11, 12^. Very few, such as Sequin^13–15^, also provides the option for users to order reference materials online. These portals mainly help users obtain the reference materials and relevant data already generated, but do not provide convenient solutions supporting the follow-up processes after access to reference materials, thus limiting the widespread application of reference materials by the community.

Continuously integrating community feedback on the use of reference materials could provide data foundation for accelerating the development of the existing quality control system. Intra- and inter-lab performance varies considerably in terms of instruments, experimental protocols, and computational pipelines^4, 16^. These differences are very important to be studied in quality control studies, but cannot be exhausted in a single study. Therefore, it is essential to integrate real-life data from communities with complementary technical strengths and complex performances. A paradigm model is the crowdsourced precisionFDA challenges, which leverages the power of community participants to identify the QC tools with high accuracy and robustness^17^, and to upgrade benchmarks for easy- and difficult-to-map genomics regions^18^, etc. This exemplary model deserves to be extended to more dimensions with other types of omic studies to help researchers gain the knowledge and resources to ensure data quality and thus improve the reliability of omics-based biological discoveries.

In this context, we developed the Quartet Data Portal around the Quartet Project. The Quartet Project was launched for quality control of multiomic profiling based on the large quantities of multiomic reference materials derived from immortalized B-lymphoblastoid cell lines of a monozygotic twin family. See accompanying papers on the overall project findings^19^, genomics^20^, transcriptomics^21^, proteomics^22^, metabolomics^23^, and batch-effect monitoring and correction^24^ with the Quartet multiomics reference materials. With the community-wide efforts, extensive datasets across platforms, labs, protocols, and batches were generated for the multiomic characterization of the reference materials. The Quartet Project team has developed the corresponding reference datasets, QC metrics, and analysis tools for genomics, transcriptomics, proteomics, and metabolomics to accompany the reference materials, resulting in a comprehensive quality control system. The Quartet Data Portal is a central hub that integrates all these resources and is dedicated to promoting the use of reference materials and to continuously upgrade the Quartet quality control system. Functions provided include channels for requesting multiomic reference materials, tools for obtaining multi-level data, interactive visualization for exploring reference datasets, and online applications for assessing user-submitted data quality. The portal is compliant with the FAIR (Findability, Accessibility, Interoperability, and Reusability) principles and is aimed for advancing scientific data management and community sharing efforts^25^.

## Results

### Overview of the Quartet Data Portal

The Quartet Data Portal integrates rich resources of the Quartet Project and consists of four core components (**Fig. 1a**): (1) Reference Materials. A unique online channel is provided for the public to request reference materials. Essential information on reference DNAs, RNAs, proteins, and metabolites is displayed in this module. (2) Multiomic Data. A data hub for accessing multi-level omic data, which involves metadata, raw datasets, intermediate datasets, and profiles. (3) Quality Assessment. Analytical tools are developed to assess the quality of user-submitted data and to generate quality assessment reports. (4) Reference Datasets. This module contains the reference datasets of high-confidence small variants (SNVs and Indels), structural variants (SVs), RNAs, proteins, and metabolites, as well as interactive visualization tools for quick understanding and exploration.

**Fig. 1.**
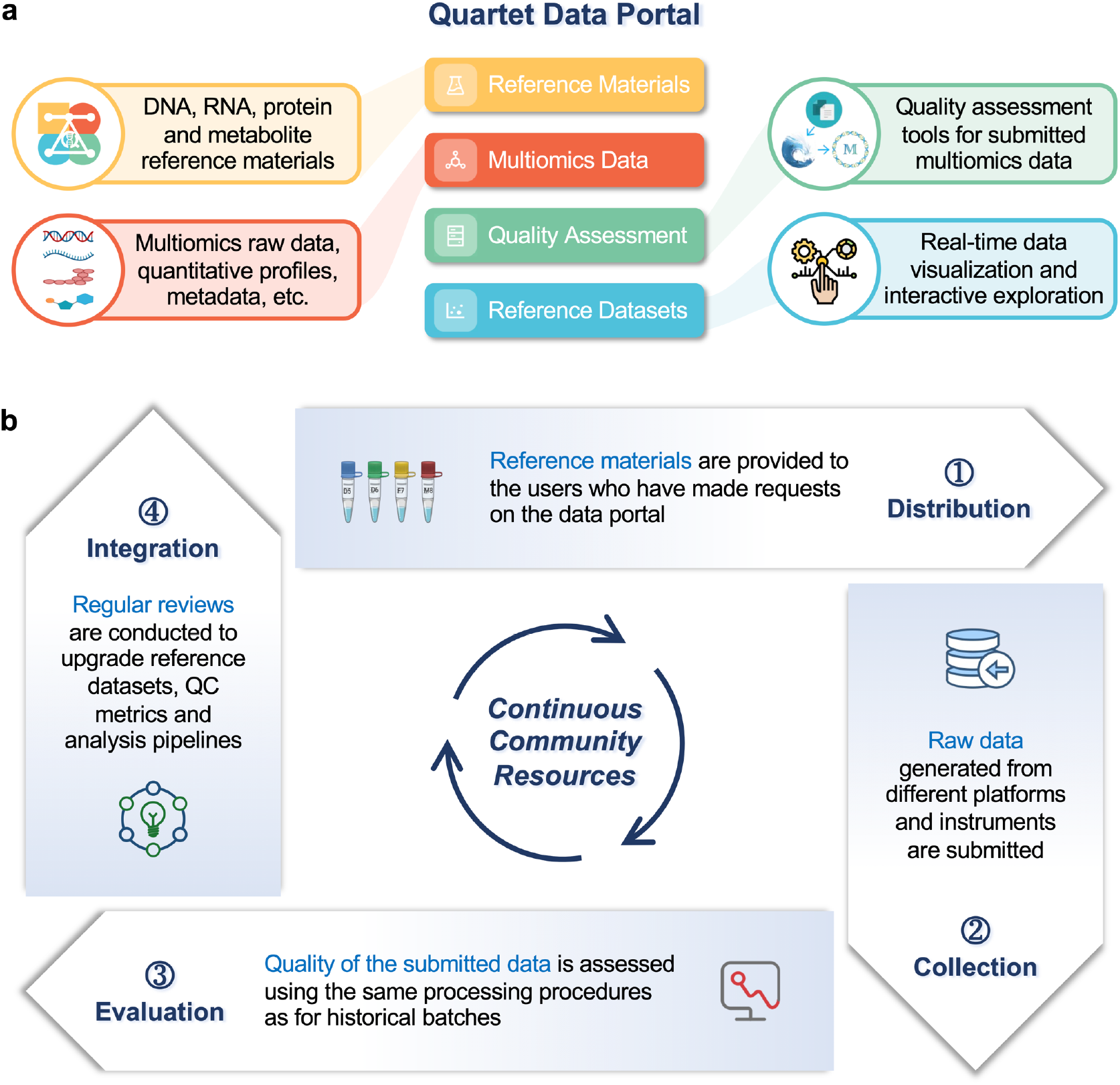
Overview of the Quartet Data Portal. **a**, Reference materials, multiomic data, quality assessment, and reference datasets constitute the core of the Quartet Data Portal, a user-friendly web-server for requesting samples, accessing to multiomic data, evaluating quality of submitted data, and visualizing reference datasets. **b**, The Quartet Data Portal uses a “distribution-collection-evaluation-integration” closed-loop workflow. Continuous requests for reference materials by the community will generate large amounts of data from the Quartet reference samples under different platforms and labs. This helps improve the quality control metrics and analysis pipelines, as well as to gradually improve the reliability and application scope of the reference materials and datasets.

### The “distribution-collection-evaluation-integration” model supports the continuous evolution of technologies

Users who have applied for the reference materials are encouraged to upload raw data from sequencing or mass spectrometry back to the Quartet platform for analysis. At this point, the Quartet Data Portal enables an effective interaction with the community, forming a closed-loop “distributioncollection-evaluation-integration” workflow (**Fig. 1b**). This offers mutual benefit for both the users and the Quartet platform. Users can obtain a comprehensive quality assessment report and all relevant data from the analysis process, whereas the Quartet team can further improve the overall quality control system by leveraging newly submitted data with a stringent process of review and integration.

The data collected from the community by the Quartet Data Portal include, but are not limited to, (1) constantly growing and manually curated multi-level data of genomics, transcriptomics, proteomics, and metabolomics, generated across platforms, labs, protocols, and batches; (2) regularly upgraded multiomic reference datasets with accompanying QC metrics based on the above data sources, and (3) version-controlled data analysis pipelines for the quality assessment of multiomic data.

### Multiomic reference materials and multi-level data resources are publicly available

The resources of the Quartet Data Portal cover the whole process of multiomic data generation and data analysis in the Quartet Project (**Fig. 2a**). The Quartet reference materials are extracted from immortalized B-lymphoblastoid cell lines (LCLs), which were established from the peripheral blood samples of four members of a family Quartet including father (F7), mother (M8), and monozygotic twin daughters (D5 and D6). Thousands of vials of reference DNAs, RNAs, proteins and metabolites that have already been verified for homogeneity and stability are stored in the −80°C freezers (**Fig. 2b**).

**Fig. 2.**
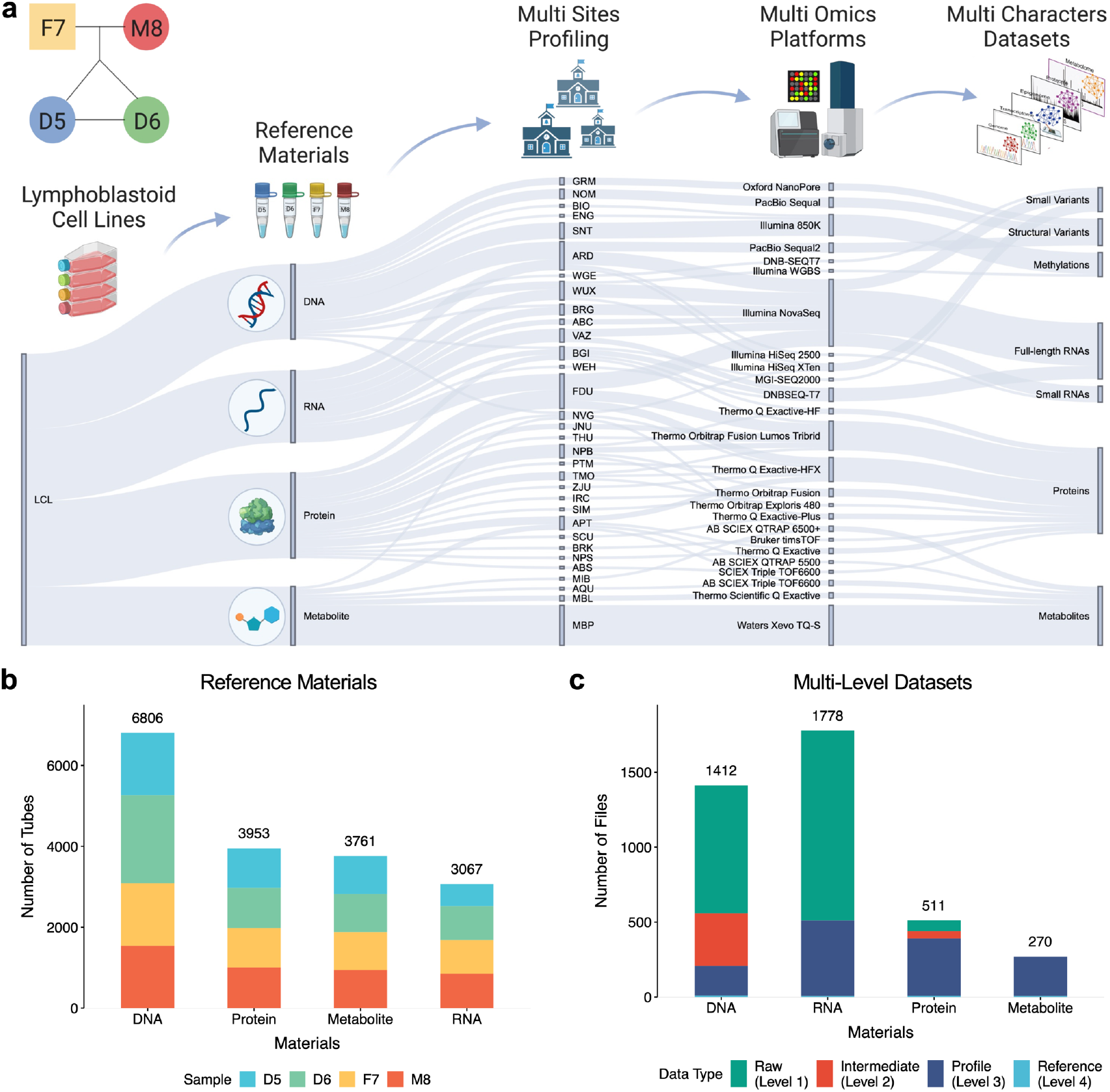
Available reference materials and data resources. **a**, The Sankey diagram displays a general view of the Quartet Project supported by the Quartet Data Portal. **b**, Bar plot of the number of available Quartet multiomic reference materials. **c**, Bar plot of the number of files for the raw datasets (level 1), the intermediate datasets (level 2), the profiles (level 3), and the reference datasets (level 4).

Large quantities of data cross six omic types, 25 platforms, and 32 labs are collected and available via the Quartet Data Portal. Genomic data were measured on four short-read (Illumina HiSeq and NovaSeq, BGI MGISEQ-2000 and DNBSEQ-T7) and three long-read (Oxford Nanopore Technologies (ONT), Pacific Biosciences (PacBio) Sequel and Sequel II) sequencing platforms at eight labs. Methylomic data included Illumina EPIC (850K) array data and whole-genome bisulfite sequencing (WGBS) data measured at four labs. Full-length RNA sequencing data were generated on MGI DNBSEQ-T7 and Illumina NovaSeq using poly(A) selection or RiboZero library preparation protocols at eight labs. Small RNA sequencing data were generated on Illumina NovaSeq and HiSeq 2500 at four labs. Peptides were measured on nine LC-MS/MS based proteomics platforms (Thermo Scientific Q Exactive, Q Exactive-HF, Q Exactive-HFX, Q Exactive-Plus, Orbitrap Fusion Lumos Tribrid, Orbitrap Fusion, Orbitrap Exploris 480, Bruker timsTOF, and SCIEX Triple TOF6600) at 16 labs. Metabolites were measured on five LC-MS/MS based metabolomics platforms (Thermo Scientific Q Exactive, SCIEX Triple TOF6600, QTRAP 6500+, QTRAP 5500, and Xevo TQ-S) at six labs (**Fig. 2a**).

Over all, the Quartet Data Portal provides more than 100 TB of 3917 multi-level data files including raw data (level 1), intermediate data (level 2), profiles (level 3), and reference datasets (level 4) that have been managed hierarchically (**Fig. 2c**). All levels of genomic data are available; intermediate files (Binary Alignment Map files) are not retained for transcriptomic data; and for metabolomics, only profiles and reference datasets are provided. In addition, metadata involved in the whole process, from study design to the final step of data analysis, are documented and available. In the first release, a total of 5,373,058 small variants (SNVs and Indels), 19,129 structural variants (SVs), 15,372 full-length mRNAs, 3,197 proteins (annotated from peptides), and 82 metabolites are contained in the reference datasets.

Users can obtain the characteristics of the reference materials and apply for the samples through an online channel (**Fig. 3a**). After approval by the Quartet team, the reference materials will be distributed to the users. Multi-level datasets and structured metadata can be obtained in a data hub which provides the browser, search, and download functions. As shown in the Pie charts of files that meet the filtering conditions and search criteria at the same time (**Fig. 3b**), a faceted search interface allows the filtration according to omic type, data category, data format, platform, protocol, etc. Finally, the flat files for the retrieved datasets and the corresponding metadata can be downloaded. Here, we structured the metadata across the whole experimental process, and designed the nodes and their properties according to the NCI Thesaurus (NCIt) standard^26^. The properties of each node, as well as its types, requirements, and descriptions are predefined. Importantly, users who have obtained the Quartet reference materials can fill in the metadata template and return it along with the multiomic data generated in their own institutions. Detailed procedures for data acquisition are described in this website: https://docs.chinese-quartet.org/about/policies/.

**Fig. 3.**
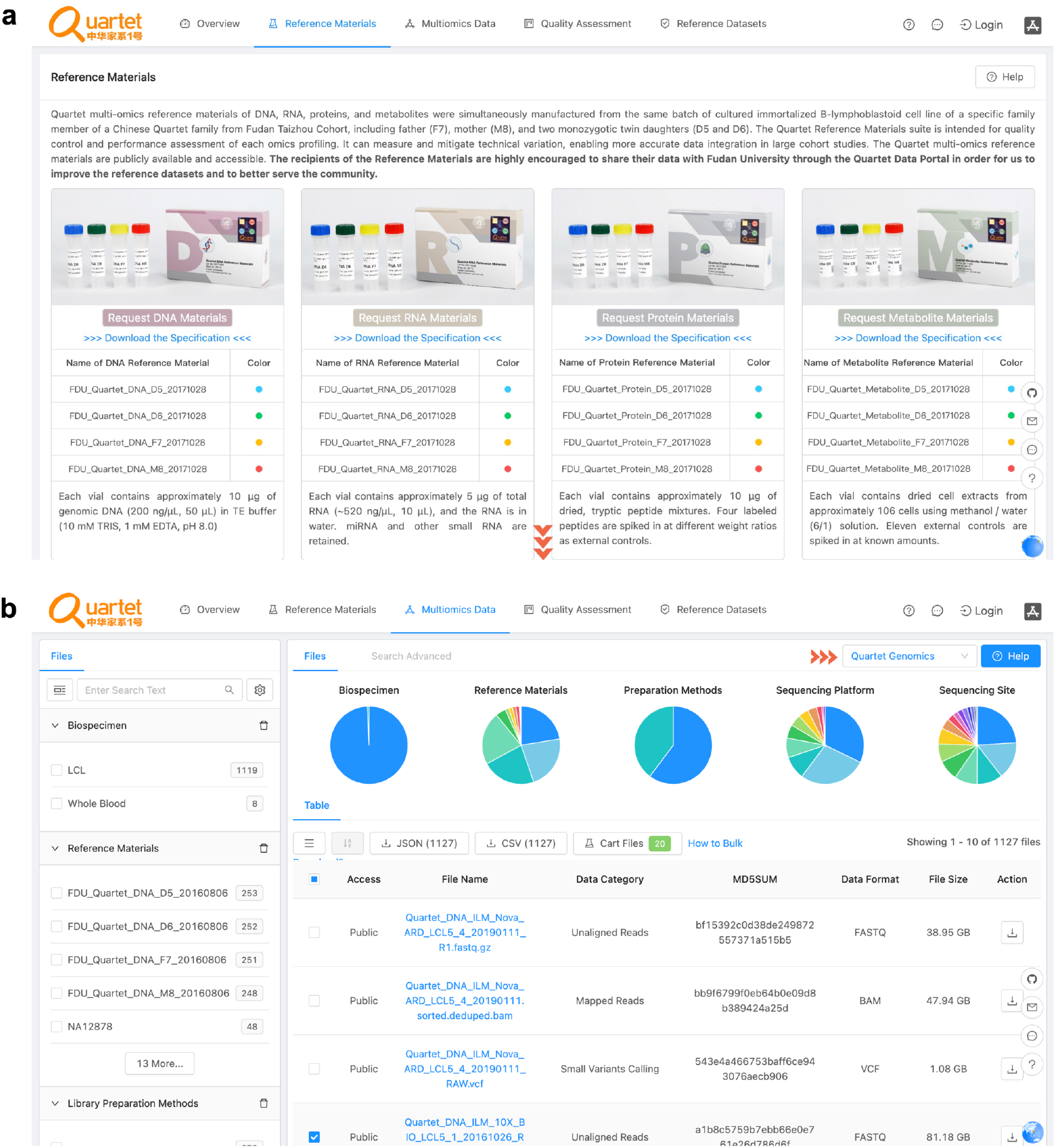
Interfaces for requesting reference materials and reference datasets from the Quartet Data Portal. **a**, Interface for requesting Quartet multiomic reference materials. **b**, Interface for requesting Quartet multiomic datasets.

### Interactive visualization enables instant exploration of the reference datasets

Understanding the characteristics of the reference datasets is essential but needs to be made relatively intuitive for users who want to utilize the Quartet quality control system. This is especially challenging for visualizing the genomics and transcriptomics data in multiple dimensions. In this regard, interactive visualization tools were developed to assist users in quickly exploring the reference datasets.

It features the following three functionalities. The first function is to perform a real-time query for the expression of specific genes, proteins, and metabolites (**Fig. 4a**). This function enables users to retrieve the expression level of the query objects under different conditions, e.g., samples, labs, protocols, instruments, etc. The second function is to integrate pre-processing methods for real-time calculation and visualization (**Fig. 4b**). This function allows users to select different batch combinations and corresponding methods for the correction of batch effects. The Principal Component Analysis (PCA) figures plotted in real-time can help the user choose the most appropriate ones. Finally, it allows the users to select the visualization module around the perspective that QC metrics focus on. As shown in **Fig. 4c**, which is a partial example of the visualization module for transcriptomics, users can understand the reference dataset better from different perspectives by using different display of interactive visualizations.

**Fig. 4.**
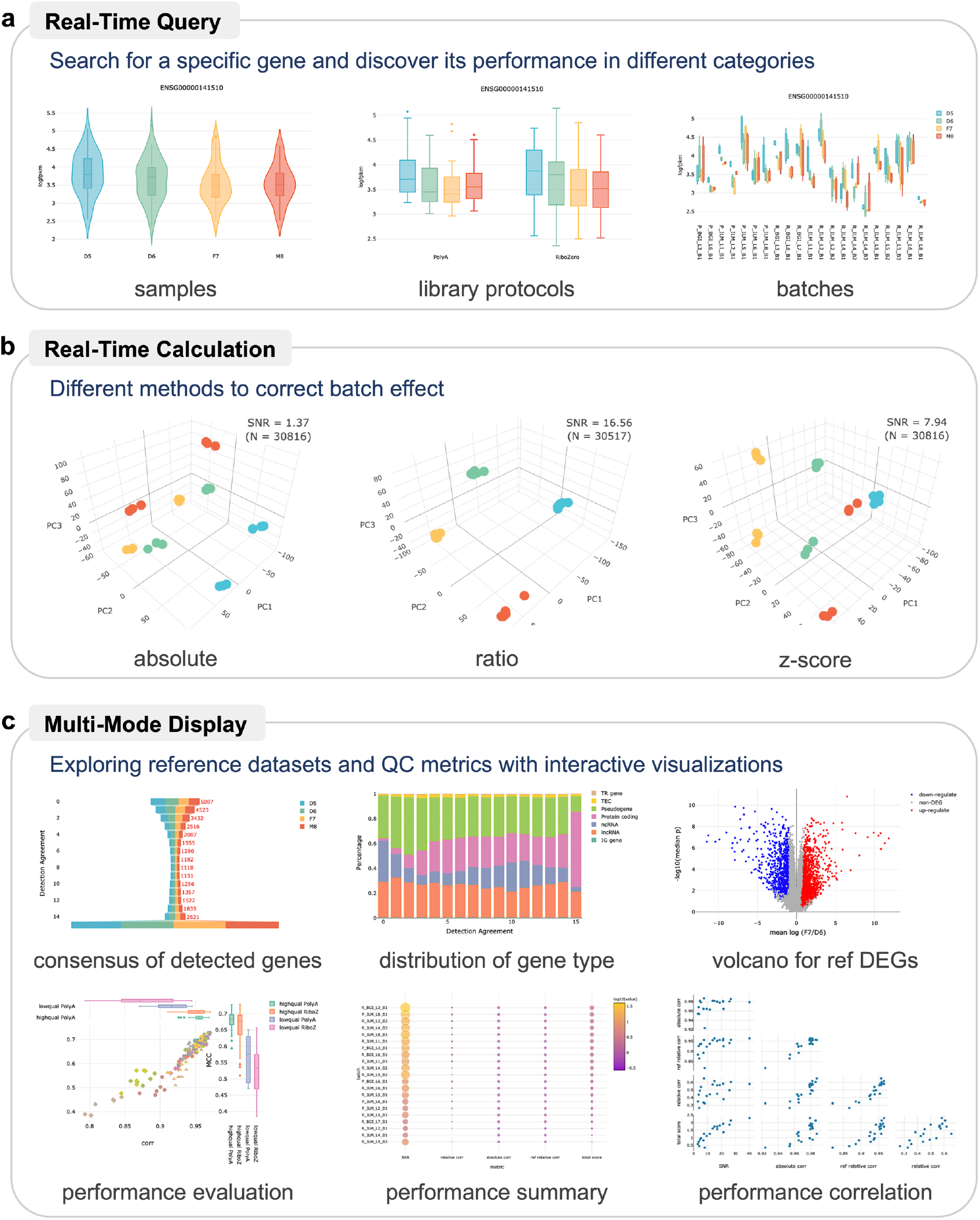
Online interactive visualization for instant exploration of the reference datasets. **a**, Real-time query to obtain the expression status of a specific gene. **b**, Realtime calculations are performed when different pre-processing methods are selected. **c**, Multi-mode display approach helps users explore reference datasets from multiple perspectives.

### User-generated multiomic data can be submitted online for real-time quality assessment

Online quality assessment has been built as part of applications in the Quartet Project quality control system. Users can upload their in-house omics data generated with the Quartet reference samples, select the specific pipeline and parameters, and obtain the analysis results and QC results (**Fig. 5a**). The Quartet Data Portal will provide private storage of the above files for a specific period of time, meaning that users can query, download, and delete their analysis results themselves during this cycle.

**Fig. 5.**
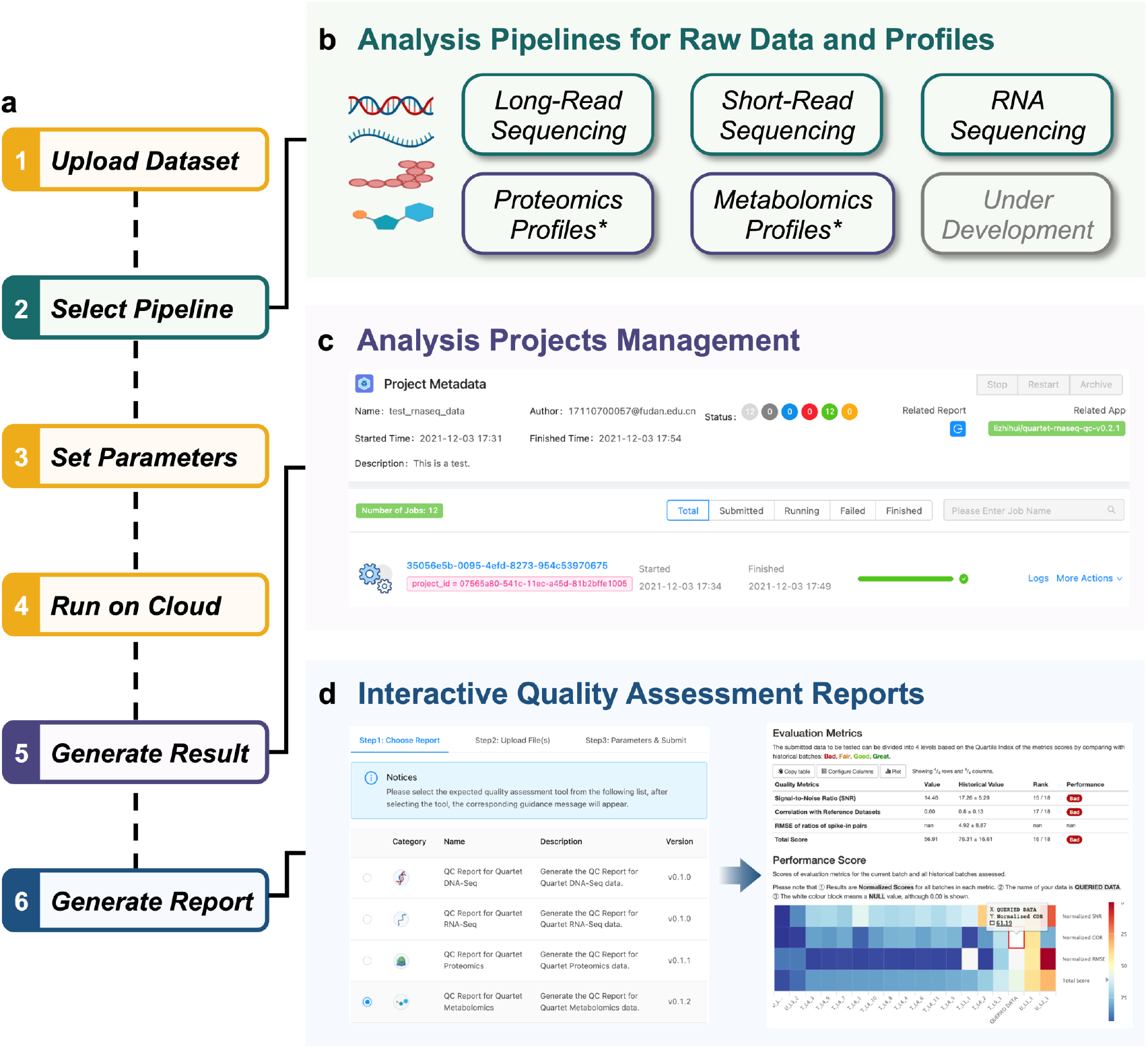
Quality assessment for user-submitted omic data. **a**, A typical workflow for quality assessment of omic data on the Quartet Data Portal. **b**, Analytical functionalities are organized as reproducible pipelines for long-read sequencing, short-read sequencing, RNA sequencing, proteomics profiles, and metabolomics profiles. **c**, Historical analysis projects are managed by the data portal. The basic information, metadata, results of tasks, etc., are all available. **d**, Snapshots of creating an interactive quality assessment report and the generated report.

Currently, the portal provides the bioinformatic analysis pipelines for long-read sequencing, short-read sequencing, RNA sequencing, proteomics profiles, and metabolomics profiles (**Fig. 5b**). These applications start with raw data of short-read sequencing, long-read sequencing and RNA sequencing and obtain variant calling files (VCF) or gene expression profiles. All historical analysis projects are available in individual accounts, and the status of calculations and related results for each sample in each individual project can be downloaded (**Fig. 5c**).

For proteomics and metabolomics, the development of full-flow applications starting from raw data is limited by the mass spectrometry technology itself^12^. Unlike transcriptomics, the subsequent analysis of the raw mass spectrometry data is platform-dependent, which is reflected in different parsing software, annotation databases, etc. In this case, we provide QC tools to support quality assessment from the profile data onwards, i.e., directly start from step 6 of **Fig. 5a**.

The last and most important step is the generation of the quality assessment report (**Fig. 5d**). Users who analyze genomic or transcriptomic data can directly select the corresponding project ID, while users who analyze proteomic or metabolomic data need to upload profiles and set parameters. This session will further calculate the scores of the submitted data in the QC metrics of the corresponding omics. The ranking of the submitted dataset will be provided by comparing its performance with that of representative historical batches integrated in the Quartet Project. Finally, the portal provides an overall evaluation based on the ranking quartiles, which are specifically classified into four categories, i.e., great, good, fair, and bad.

### Application scenarios implemented with the Quartet Data Portal

A general application process involves the first three steps in the closed-loop process mentioned earlier (**Fig. 6a**). 1) Users request the Quartet reference materials and then perform experiments and generate data from these samples individually, or in each batch along with biological study samples and generate data. 2) The raw data from the Quartet reference materials can be submitted back to the data portal and evaluated for quality using the platform’s analysis pipelines. 3) The submitted data will be scored, ranked, and given an overall performance category based on the QC metrics and Quartet reference datasets.

**Fig. 6.**
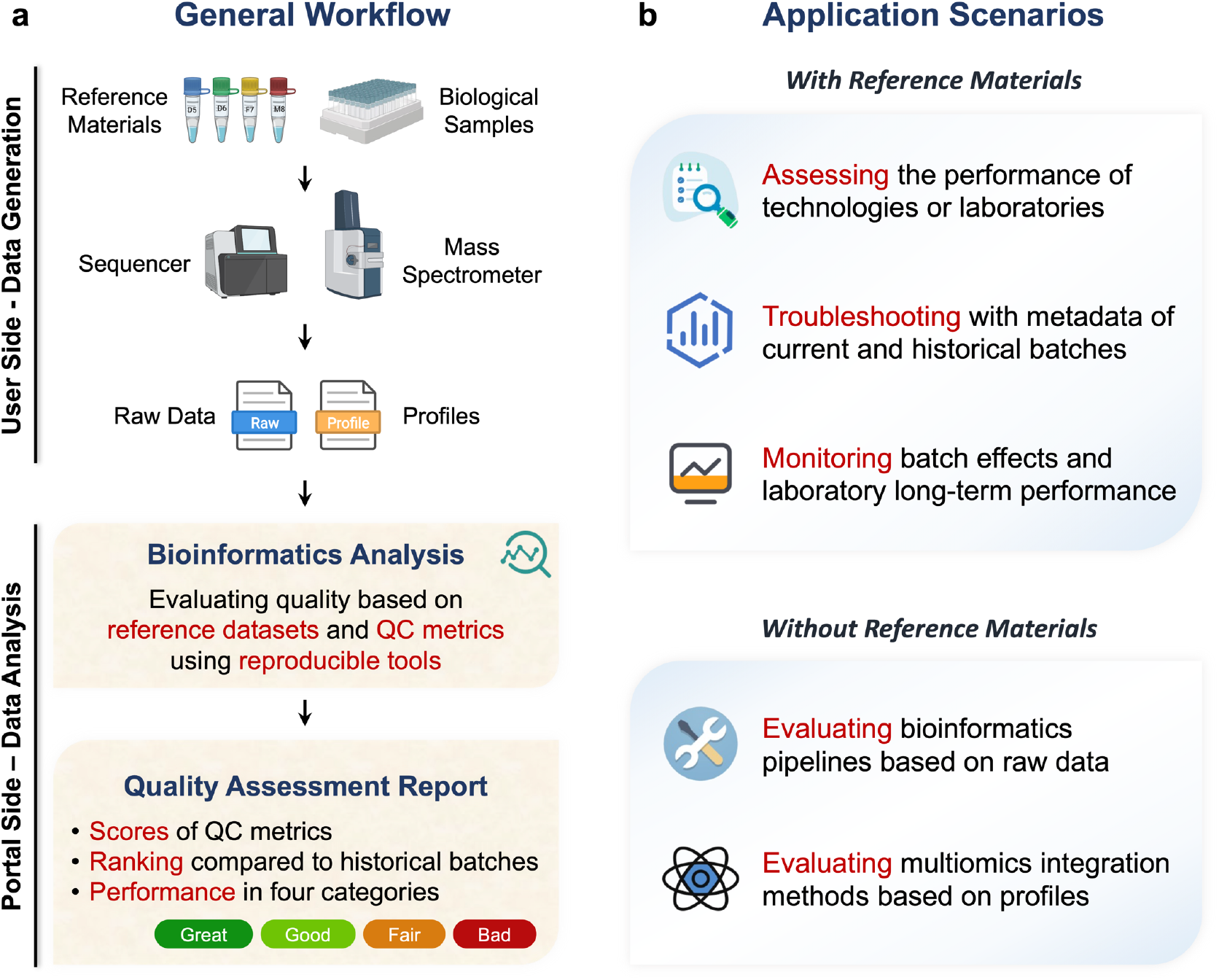
Application scenarios of the Quartet Data Portal. **a,** General workflow of the main applications. The user performs experiments on reference samples and biological samples within the same batch. And after generating sequencing or mass spectrometry data, the user submits the resulting data to the Quartet Data Portal. The system will run a reproducible processing procedure to generate a comprehensive quality assessment report for the user. **b,** Application scenarios with or without reference materials.

Following the above process, users can achieve the following, as shown in **Fig. 6b**. First, the users can use the Quartet reference materials to evaluate the quality of their inhouse data generation steps. In particular, reference materials can play an important role in assessing the performance of new technology platforms and reagents, or the qualifications of service labs or operators. Secondly, in the multi-batch experiments, reference materials from each batch can be used to monitor batch effects and long-term lab performances. Besides, users can correct the batch effects based on the relative expression of Quartet samples (detailed in accompanying articles). Thirdly, metadata of historical batches of the Quartet Project is helpful for troubleshooting. Sequencing and mass spectrometry experiments are complex and composed of multiple steps, but metadata records detailed experimental information about every step. Influencing factors can be eliminated case by case by comparing with other studies.

In addition, other users who have not applied for the Quartet reference materials can also make more in-depth use of the resources provided by Quartet Data Portal. Here we provide two scenarios. On the one hand, large amounts of raw data of the Quartet Project can be used to evaluate the performance of various bioinformatics pipelines. Specifically, the same sequencing data can be analyzed by using different pipelines and their performances can be evaluated based on the comparison of the obtained profiles with the reference datasets. On the other hand, multiomic profiles and the Quartet study design with built-in biological truth can be used to explore multiomic integration issues, e.g., performance evaluation of integration tools.

## Discussion

The Quartet Data Portal integrates the unique resources and results from the Quartet Project with various tools, e.g., interactive visualization and QC, to facilitate the application of the quality control system based on the Quartet reference materials in the community. Currently, the portal involves multiple steps such as data storage, data analysis, and quality report. Guided by the FAIR principles^25^, multi-level data related to the Quartet Project are managed and published uniformly by the platform in a version-controlled fashion. To date, multiple studies based on data published by the Quartet Data Portal have been successfully conducted^27, 28^. Besides, a complete process has been established for the quality assessment of sequencing data and mass spectrometry profiles. However, the online analysis of proteomics and metabolomics raw datasets to expression profiles is currently not yet available. Users can only upload profiling results in the specified format. In the future, we will further improve the QC tools for proteomic and metabolomic analysis based on more community data and more advanced analysis software.

With the widespread use of the Quartet reference materials by the community, we will inevitably face the challenge of integrating data from different sources and the continuous evolution of the QC system integrated in the Quartet Data Portal^29^. The importance of reproducible data analysis pipelines and structured metadata formats for the community-wide multiomic research is well-acknowledged^30–32^. Consequently, we have built this Portal to address these unmet needs. In terms of metadata specification, we have structured the metadata of the whole experiment and analysis processes and defined the fields with the knowledge of ontology. This helps users interpret the data correctly and also makes comparisons with other studies more straightforward and meaningful. To ensure the computational reproducibility, analysis pipelines are strictly versioned, and the software within the pipelines are packaged using docker images^33^. The QC tools that users can choose from are also consistent with the version of the reference datasets and QC metrics.

The “distribution-collection-evaluation-integration” model implemented in the portal allows more researchers to be truly involved in the quality control of multiomic studies. However, there are still some challenges that need to be addressed step-by-step in the future. First, the current release of multi-level datasets requires the manual collation by the Quartet team on a regular basis. In the future, we plan to develop automated execution processes as well as more granular provisions to avoid the potential delay and bias of manual processing. In addition, a wider variety of reproducible pipelines and analysis models (e.g., local standalone software) are envisioned to help researchers perform more personalized analyses.

In summary, we have made an attempt in promoting the multiomics research community to work together to solve quality problems. Our intention is to integrate community-generated data while sharing the public with Quartet reference materials. The “distribution-collection-evaluation-integration” closed-loop model drives the evolution of reference datasets, QC metrics, and QC tools for small variants, structural variants, mRNAs, proteins, and metabolites. We believe that the Quartet Data Portal can be useful for multiomic studies, helping raise awareness of quality control among researchers in the community and laying a solid foundation for more reliable biological discoveries.

## Methods

The construction of the Quartet Data Portal consists of four main parts: multiomic data management module, data analysis and quality assessment module, quality report module, and visualization dashboard module.

### Multiomic Data Management Module

The Quartet Data Portal has a set of solutions that support flexible customization and expansion of metadata, and can reflect the structural relationship between metadata and support version upgrades and evolution. The current metadata solution includes project, donor, sample, reference material, library, sequencing, datafile and other entity information to track the details of the entire process of data generation and analysis. This module relies on the NoSQL databases including MongoDB and Nebula Graph DB, to handle the storage of a large amounts of semi-structured data, and object storage, i.e., S3, MinIO, Alibaba Cloud Object Storage Service (OSS), to realize the storage of a large number of omics data files.

The verification of the Quartet metadata is a key step for effective data management. All metadata is included in the unified management system and needs to follow strict data types, limit value ranges, and check for validity of values, etc., while supporting metadata model extension and version replacement. Therefore, we have defined a set of “Data Package” specifications to implement the constraints and verification of metadata. It is mainly composed of a set of specific directory structures and several CSV files. Each CSV file corresponds to a model description file based on JSON schema to complete the verification of the corresponding data structure and contents.

### Data Analysis and Quality Assessment Module

Multiomic data usually require a relatively large storage and computation capacities. For quality control of the raw data, computing resources are required, computing time is long, and quality control pipelines related to different omics data are different. The Quartet Data Portal requires a set of computing systems dedicated to multiomic data analysis and flexible definition of quality control pipelines to build quality assessment modules. Therefore, we have defined a set of specifications to realize the encapsulation and definition of the pipelines, which is mainly based on Workflow Definition Language (WDL), template language and catalog specifications. In order to meet the needs of cross-platform scheduling computing tasks, we implemented a computing system that supports custom application specifications based on the Cromwell scheduling engine. It is combined with a series of omics data quality control applications to complete the quality assessment of multiomic data.

### Quality Report Module

There are many quality evaluation indicators, and the content of the quality control report of different types of omic data is different. The Quartet Data Portal requires a set of quality assessment report modules that support custom report contents and styles and can be used to interactively explore results. Therefore, we use Clojure language to implement a set of quality assessment report modules that support plug-in mechanism, and all report plug-ins are built based on MultiQC^34^ and Plotly (https://github.com/plotly/plotly.py). Report plug-ins can be added and deleted flexibly to complete the generation and display of corresponding quality assessment reports.

### Visualization Dashboard Module

The visualization dashboard is developed in the R language (version v3.5.1) (https://www.r-project.org). The shiny v1.2.0 and shinydashboard v0.7.1 are used to deploy on a self-managed webserver based on the Shiny Server (https://www.rstudio.com/products/shiny/shiny-server/). The widgets of the webserver are developed by shinyWidgets v0.4.8. The analysis tables are processed by data.table v1.12.8 and feather v0.3.5 due to their high efficiency, and data are manipulated using the plyr v1.8.4, dplyr v1.0.2 and purrr v0.3.2. The visualization module is developed using the R package plotly v4.9.2.1, which includes scatter, box, bar, violin, heatmap, funnel and splom types. Except for the heatmap and dendrogram, the R package heatmaply v1.1.1 is used. The other section of the genome visualization module uses R package plotly v4.9.2.1 and ggplot2 v3.3.2, which includes scatter, line, box and bar types. The theme of dashboard is developed through dashboardthemes v1.1.3, and the theme of ggplot2 is mainly through ggthemes v4.2.0. The colors of the plots are set using RcolorBrewer v1.1-2. The data values are mapped to the graph using lattice v0.20-35 and scales v1.1.1. Specifically, the renv v0.11.0 R package is invoked to bring project-local R dependency management to the project.

## Data availability

All data are available in the Quartet Data Portal, which can be accessed at https://chinese-quartet.org. They can also be accessed from the Genome Sequence Archive (GSA) of the National Genomics Data Center of China with BioProject ID of PRJCA007703. The documentation is available here: https://docs.chinese-quartet.org.

## Code availability

The source code can be obtained through https://github.com/chinese-quartet.

## Disclaimer

The content is solely the responsibility of the authors and does not necessarily represent the official views of the US Food and Drug Administration.

## Acknowledgments

This study was supported in part by National Key R&D Project of China (2018YFE0201603 and 2018YFE0201600), the National Natural Science Foundation of China (31720103909 and 32170657), Shanghai Municipal Science and Technology Major Project (2017SHZDZX01), State Key Laboratory of Genetic Engineering (SKLGE-2117), and the 111 Project (B13016). We are grateful to visitors to the Quartet Data Portal for their feedback that helped us improve the functionalities and user-friendliness of the portal. Some of the illustrations in this paper were created with BioRender.com.

## Author contributions

Y.Z., W.X., L.S., H.H., W.T., J.Y., and Y.L. conceived and designed the study. J.Y., Y.L., and Q.W.C. designed the metadata specifications for the data module. L.R., N.Z, Q.C.C., Z.L., and Y.Y. curated the multiomic data. J.Y. and J.S. integrated all analysis applications and reporting plugins contributed by L.R. and Y.L. (Genomics), Z.L. and J.S. (Transcriptomics), Q.C.C. and Y.L. (Proteomics), N.Z. and Y.L. (Metabolomics). J.S. and Y.S. developed visualization plugins for genomics and transcriptomics respectively, and J.Y. integrated them into the portal. Q.C.C., Y.L., S.Y., A.S., J.Y., S.Y. and others contributed to the testing and public release. Y.L., Y.Z. and J.Y. drafted the manuscript; W.X., L.S., H.H., W.T., and A.S. revised it. All authors reviewed and approved the manuscript.

## References

1. Gargis, A.S. et al. Assuring the quality of next-generation sequencing in clinical laboratory practice. Nat. Biotechnol. 30, 1033–1036 (2012).

2. Hardwick, S.A., Deveson, I.W. & Mercer, T.R. Reference standards for next-generation sequencing. Nat. Rev. Genet. 18, 473–484 (2017).

3. Salit, M. & Woodcock, J. MAQC and the era of genomic medicine. Nat. Biotechnol. 39, 1066–1067 (2021).

4. Shi, L. et al. The MicroArray Quality Control (MAQC) project shows inter-and intraplatform reproducibility of gene expression measurements. Nat. Biotechnol. 24, 1151 (2006).

5. Su, Z. et al. A comprehensive assessment of RNA-seq accuracy, reproducibility and information content by the Sequencing Quality Control Consortium. Nat. Biotechnol. 32, 903–914 (2014).

6. Fang, L.T. et al. Establishing community reference samples, data and call sets for benchmarking cancer mutation detection using whole-genome sequencing. Nat. Biotechnol. 39, 1151–1160 (2021).

7. Xiao, W. et al. Toward best practice in cancer mutation detection with whole-genome and whole-exome sequencing. Nat. Biotechnol. 39, 1141–1150 (2021).

8. Zook, J.M. et al. Integrating human sequence data sets provides a resource of benchmark SNP and indel genotype calls. Nat. Biotechnol. 32, 246–251 (2014).

9. Zook, J.M. et al. An open resource for accurately benchmarking small variant and reference calls. Nat. Biotechnol. 37, 561–566 (2019).

10. Orchard, S. Data standardization and sharing—the work of the HUPO-PSI. Biochimica et Biophysica Acta (BBA)-Proteins and Proteomics 1844, 82–87 (2014).

11. Beger, R.D. et al. Towards quality assurance and quality control in untargeted metabolomics studies. Metabolomics 15, 4 (2019).

12. Evans, A.M. et al. Dissemination and analysis of the quality assurance (QA) and quality control (QC) practices of LC–MS based untargeted metabolomics practitioners. Metabolomics 16, 1–16 (2020).

13. Deveson, I.W. et al. Representing genetic variation with synthetic DNA standards. Nat. Methods 13, 784–791 (2016).

14. Hardwick, S.A. et al. Spliced synthetic genes as internal controls in RNA sequencing experiments. Nat. Methods 13, 792–798 (2016).

15. Blackburn, J. et al. Use of synthetic DNA spike-in controls (sequins) for human genome sequencing. Nat. Protoc. 14, 2119–2151 (2019).

16. Peng, R.D. & Hicks, S.C. Reproducible research: a retrospective. Annu. Rev. Public Health 42, 79–93 (2021).

17. Yoo, S. et al. A community effort to identify and correct mislabeled samples in proteogenomic studies. Patterns 2, 100245 (2021).

18. Olson, N.D. et al. PrecisionFDA Truth Challenge V2: Calling variants from short and long reads in difficult-to-map regions. Cell Genomics 2, 100129 (2022).

19. Zheng, Y. et al. Ratio-based multiomics profiling using universal reference materials empowers data integration [Unpublished manuscript]. (2022).

20. Ren, L. et al. Quartet DNA reference materials and datasets for comprehensively evaluating germline variants calling performance [Unpublished manuscript]. (2022).

21. Yu, Y. et al. Quartet RNA reference materials and ratio-based reference datasets for reliable transcriptomic profiling [Unpublished manuscript]. (2022).

22. Tian, S. et al. Quartet protein reference materials and datasets for multi-platform assessment of label-free proteomics [Unpublished manuscript]. (2022).

23. Zhang, N. et al. Quartet metabolite reference materials and datasets for inter-laboratory reliability assessment of metabolomics studies [Unpublished manuscript]. (2022).

24. Yu, Y. et al. Correcting batch effects in large-scale multiomic studies using a referencematerial-based ratio method [Unpublished manuscript]. (2022).

25. Wilkinson, M.D. et al. The FAIR Guiding Principles for scientific data management and stewardship. Sci. Data 3, 1–9 (2016).

26. Sioutos, N. et al. NCI Thesaurus: a semantic model integrating cancer-related clinical and molecular information. J. Biomed. Inform. 40, 30–43 (2007).

27. Khayat, M.M. et al. Hidden biases in germline structural variant detection. Genome Biol. 22, 1–15 (2021).

28. Pan, B. et al. Assessing reproducibility of inherited variants detected with short-read whole genome sequencing. Genome Biol. 23, 1–26 (2022).

29. Conesa, A. & Beck, S. Making multi-omics data accessible to researchers. Sci. Data 6, 1–4 (2019).

30. Krassowski, M., Das, V., Sahu, S.K. & Misra, B.B. State of the field in multi-omics research: From computational needs to data mining and sharing. Front. Genet. 11 (2020).

31. Tarazona, S., Arzalluz-Luque, A. & Conesa, A. Undisclosed, unmet and neglected challenges in multi-omics studies. Nat. Comput. Sci., 1–8 (2021).

32. Leipzig, J., Nüst, D., Hoyt, C.T., Ram, K. & Greenberg, J. The role of metadata in reproducible computational research. Patterns 2, 100322 (2021).

33. Boettiger, C. An introduction to Docker for reproducible research. SIGOPS Oper. Syst. Rev. 49, 71–79 (2015).

34. Ewels, P., Magnusson, M., Lundin, S. & Käller, M. MultiQC: summarize analysis results for multiple tools and samples in a single report. Bioinformatics 32, 3047–3048 (2016).

